# Identifying keystone species in microbial communities using deep learning

**DOI:** 10.1101/2023.03.15.532858

**Authors:** Xu-Wen Wang, Zheng Sun, Huijue Jia, Sebastian Michel-Mata, Marco Tulio Angulo, Lei Dai, Xuesong He, Scott T. Weiss, Yang-Yu Liu

## Abstract

Previous studies suggested that microbial communities harbor keystone species whose removal can cause a dramatic shift in microbiome structure and functioning. Yet, an efficient method to systematically identify keystone species in microbial communities is still lacking. This is mainly due to our limited knowledge of microbial dynamics and the experimental and ethical difficulties of manipulating microbial communities. Here, we propose a Data-driven Keystone species Identification (DKI) framework based on deep learning to resolve this challenge. Our key idea is to implicitly learn the assembly rules of microbial communities from a particular habitat by training a deep learning model using microbiome samples collected from this habitat. The well-trained deep learning model enables us to quantify the community-specific keystoneness of each species in any microbiome sample from this habitat by conducting a thought experiment on species removal. We systematically validated this DKI framework using synthetic data generated from a classical population dynamics model in community ecology. We then applied DKI to analyze human gut, oral microbiome, soil, and coral microbiome data. We found that those taxa with high median keystoneness across different communities display strong community specificity, and many of them have been reported as keystone taxa in literature. The presented DKI framework demonstrates the power of machine learning in tackling a fundamental problem in community ecology, paving the way for the data-driven management of complex microbial communities.

## Introduction

The notion of *keystone species* has its roots in food-web ecology^1, 2^. Since Paine coined it in describing results from his pioneering field experiments in 1969, *keystone species* has been widely applied in the ecological literature. Such a broad application (and often abuse) has generated considerable confusion about what precisely a keystone species is^3^. Here we adopt the original definition by Paine, i.e., a keystone species is a species that has a *disproportionately* large effect on the *stability* of the community relative to its abundance^1, 2^. Existing methods to identify keystone species for macro ecosystems can be classified into two approaches: *experimental manipulations* and *statistical comparisions*^4^. The first approach involves the targeted removal of suspected keystone species from the community. To avoid any selection bias, this targeted removal should be done for each species in the community. This certainly raises both logistical and ethical concerns. The second approach involves comparing two communities with different abundances or presence/absence patterns of the potential keystone species. Although this approach circumvents logistical and ethical issues, it cannot draw robust conclusions due to many potential confounding factors and unmeasured abiotic variables.

Existing studies also suggest that microbial communities harbor keystone species^5–9^. Yet, the keystone species identification approaches developed for macro ecosystems are incredibly challenging to apply to large, complex microbial communities. For experimental manipulations, targeted removal of each species in a complex community is impossible with current antimicrobial techniques, not to mention the corresponding ethical concerns for host-associated microbial communities such as the human gut microbiome. As for statistical comparisons, finding two communities that differ by just one species is challenging, especially for complex host-associated microbial communities (e.g., the human gut microbiome) with very personalized compositions^10, 11^. Moreover, statistical comparisons will suffer from numerous confounding factors. For example, it has been reported that many host variables (e.g., body mass index, sex, age, geographical location, alcohol consumption frequency, bowel movement quality, and dietary intake frequency of meat/eggs, dairy, vegetables, whole grain, and salted snacks) confound gut microbiota studies of human disease^12^.

To resolve the above limitations, one may consider directly inferring a population dynamics model to predict the temporal behavior of microbial communities and then identify keystone species through numerical simulations of targeted species removal. Many methods have been developed to solve this dynamics inference problem, using temporal or steady-state data^13–15^. Yet, those methods typically require high-quality absolute abundance data, limiting their application for identifying keystone species in large, complex microbial communities. In addition, model misspecification can strongly affect the performance of those methods. We may have to use symbolic regression techniques, a machine learning method that automatically infers both the model structure and parameters from temporal data^16–20^. But again, the informativeness of the temporal data is a big concern for complex microbial communities.

A recent numerical study^21^ claimed that those highly connected species (i.e., “hubs”) in the microbial correlation network are keystone species of microbial communities^22^. We think this claim is problematic for at least two reasons. First, edges in microbial correlation networks do not represent direct ecological interactions but just statistically significant co-occurrences or mutual exclusions of species. We emphasize that the correlation or co-occurrence network is undirected and cannot be used to predict the dynamic behavior of any ecological system simply because correlation is not causation. Spurious correlations can be observed even from a simple two-species system with deterministic dynamics^23^. Second, the impact of a species’ removal naturally depends on the resident community. This underscores a fundamental challenge on the keystone species identification --- the *community specificity*, i.e., a species may be a keystone in one community but not necessarily a keystone in another community. In other words, the keystoneness of a species can be highly community-specific. Identifying keystone species based on the hubs in a microbial correlation network (constructed from a collection of microbiome samples) or any topological indices in an ecological network (with directed inter-species interactions inferred from experimental data) completely ignore this community specificity. For host-associated microbial communities such as the human gut microbiome, the community specificity means that keystone species or keystoneness can be highly personalized due to the highly personalized microbial compositions^10, 11^.

So far, few microbial taxa have been experimentally confirmed as keystones^5, 24–26^. An efficient method to systematically identify community-specific keystone taxa in complex microbial communities is still lacking^27–29^. In this work, we propose a Data-driven Keystone species Identification (DKI) framework to fill this knowledge gap. The DKI framework does not assume any particular ecological model, naturally avoiding the model misspecification issue. Moreover, the DKI framework quantifies the keystoneness of each species for each community (sample). Hence it naturally considers the community specificity of keystoneness.

We systematically validated the DKI framework using synthetic data generated from a classical population dynamics model in community ecology. We then applied it to analyze a large-scale standardized, curated human gut microbiome dataset^30^ (consisting of 1,103 species and 2,815 samples), a large-scale oral microbiome dataset^31^ (with 702 species and 1,421 samples), a large-scale soil microbiome dataset^32^ (with 298 orders and 1,160 samples), and a large-scale coral microbiome dataset^33^ (with 1,054 genera and 1,400 samples). For each dataset, we demonstrated the community specificity of keystoneness. Moreover, we found that those taxa with high median keystoneness across different communities include many keystone taxa previously reported in the literature.

## Results

### The DKI framework

Consider a particular habitat (or meta-community) that harbors a pool of 𝑁 different microbial species, denoted as Ω = {1, ⋯, 𝑁}. Suppose we have a large set of microbiome samples 𝒮 = {1, …, 𝑀} collected from this habitat. A microbiome sample 𝑠 ∈ 𝒮 can be viewed as a local community of the habitat. The species assemblage of sample 𝑠 can be represented by a binary vector 𝒛 ∈ {0,1}^*N*^with the 𝑖-th entry 𝑧_*i*_ = 1 (or 0) if species-𝑖 is present (or absent) in 𝑠. The microbial composition or taxonomic profile of this sample is characterized by a compositional vector 𝒑 ∈ Δ^*N*^ with the 𝑖-th entry 𝑝_*i*_ representing the relative abundance of species-𝑖 in 𝑠 and Δ^*N*^ is the probability simplex. We assume the collected samples roughly represent the steady states of the local communities so that they can be used to learn the assembly rules of those communities.

The DKI framework consists of two phases. In the first phase (**Fig.1a**), we *implicitly learn* the *assembly rules* of microbial communities in this habitat using a deep learning method with 𝒮 as the training data. This is achieved by learning a map from the species assemblage 𝒛 of a sample 𝑠 = (𝒛, 𝒑) to its taxonomic profile 𝒑, i.e., 𝜑: 𝒛 ↦ 𝒑. Various deep learning methods, including Multi-Layer Perceptron^34^ (MLP) or ResNet^35^ can be used to learn such map without using any population dynamic model, but with a few reasonable assumptions (e.g., the universality of microbial dynamics, steady-state samples, no true multi-stability, and enough training samples) to ensure the problem is mathematically well-defined. Here, based on our previous work^36^, we developed cNODE2 (composition Neural Ordinary Differential Equation version 2.0) to learn the map 𝜑 (see **SI Sec.2** for details). Learning this map 𝜑 will enable us to predict what will happen to the taxonomic profile of a local community upon any species removal.

**Figure 1:**
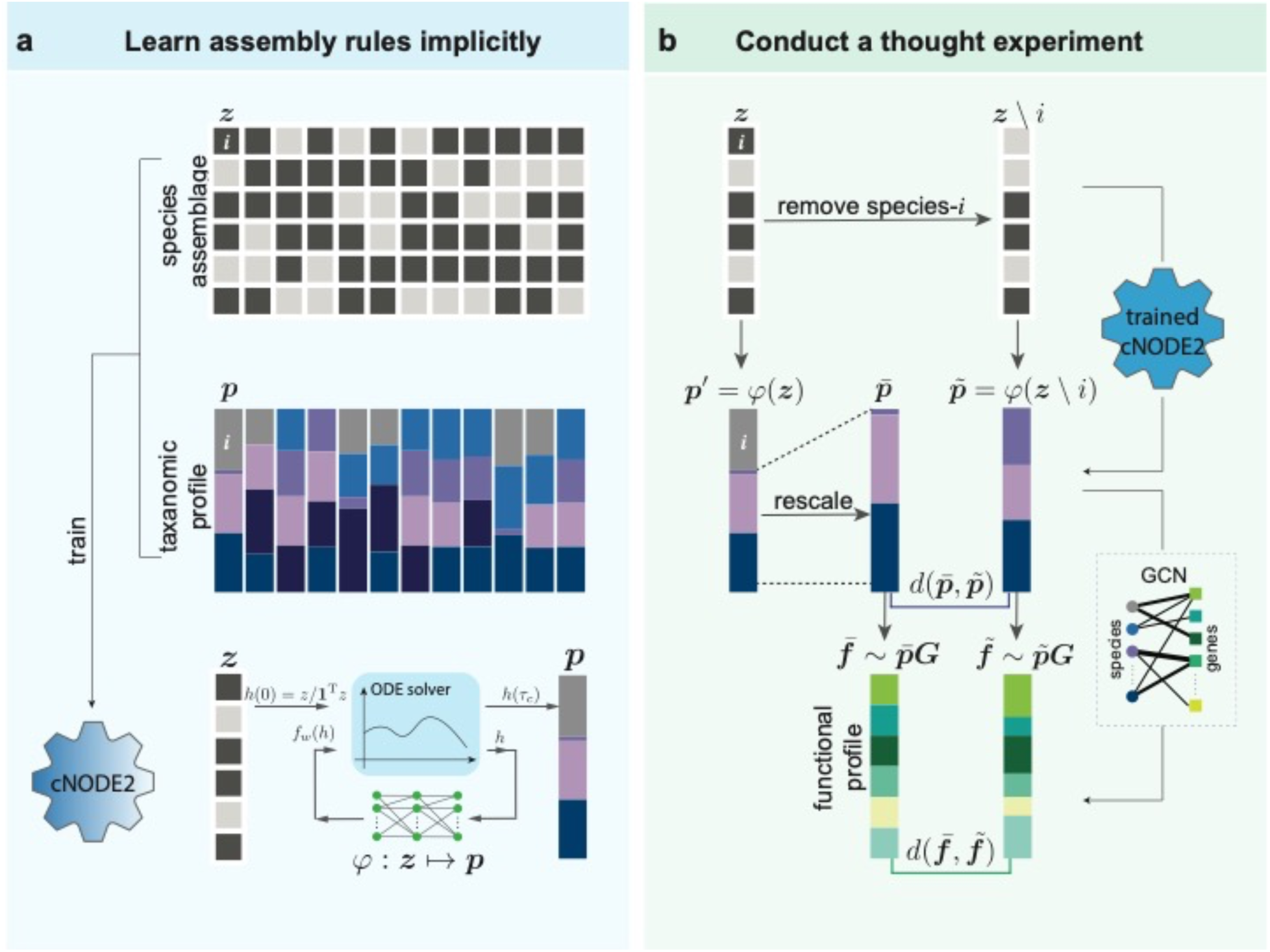
Workflow of the Data-driven Keystone species Identification (DKI) framework. a, The species assemblage of a microbiome sample 𝑠 is represented by a binary vector 𝒛 ∈ {0,1}^*N*^, where its 𝑖-th entry satisfies 𝑧_*i*_ = 1 (𝑧_*i*_ = 0) if species-𝑖 is present (or absent) in this sample. The microbial composition of this sample is characterized by a vector 𝒑 ∈ Δ^*N*^, where its 𝑖-th entry 𝑝_*i*_is the relative abundance of species-𝑖 in this sample and Δ^*N*^ is the probability simplex. A deep learning model (cNODE2) is trained to learn the map: 𝒛 ∈ {0,1}^*N*^ ↦ 𝒑 ∈ Δ^*N*^. **b,** We conduct a *thought removal experiment* of species-𝑖. In particular, for the community 𝑠 = (𝒛, 𝒑) with species collection 𝒛 and microbial composition 𝒑, we remove species-𝑖 from 𝒛 to form a new species collection 𝒛̃ = 𝒛\𝑖. Then, for the new species collection 𝒛̃, we use cNODE2 to predict its new composition 𝒑= = 𝜑(𝒛̃). To quantify the impact of species-𝑖’s removal, we need to compare the new composition 𝒑= with a null composition 𝒑> in the absence of species-𝑖 (obtained by assuming species-𝑖’s removal will not affect other species’ abundances at all). The structural impact of species-𝑖’s removal on the community 𝑠 = (𝒛, 𝒑) can be defined as the distance or dissimilarity between 𝒑= and 𝒑>, i.e., 𝑑(𝒑=, 𝒑>). Similarly, the functional impact of species-𝑖’s removal on the community 𝑠 = (𝒛, 𝒑) can be defined as the distance or dissimilarity between 𝒇̃ and 𝒇̅, i.e., 𝑑(𝒇̃, 𝒇̅). Here, the functional profile 𝒇̃ (or 𝒇̅) can be computed by multiplying 𝒑̃ (or 𝒑̅) with the incidence matrix of the genomic content network (GCN), respectively.

In the second phase (**Fig.1b**), to quantify the community-specific keystoneness of species-𝑖 in a local community or microbiome sample 𝑠, we *conduct a thought experiment* of removing species-𝑖 from 𝑠 and use cNODE2 to compute the impact of species-𝑖 ’s removal on 𝑠. In particular, for 𝑠 = (𝒛, 𝒑) with species collection 𝒛 and microbial composition 𝒑, we remove species 𝑖 from 𝒛 to form a new species collection 𝒛̃ = 𝒛\𝑖. Then, for the new species collection 𝒛̃, we use cNODE2 to predict its composition 𝒑̃ = = 𝜑(𝒛̃). To quantify the impact of species-𝑖 ’s removal, we need to compare the new composition 𝒑̃ with a null composition 𝒑̅. Here the null composition is obtained by assuming that species-𝑖’s removal will not affect other species. In case the deep-learning method can learn the map 𝜑 perfectly and predict 𝒑̃ without any error, we can directly calculate the null composition 𝒑̅ by simply setting 𝑝̅_*i*_ = 0, and renormalizing the relative abundances of the remaining species, i.e., 𝑝̅_*j*_ = 𝑝_*j*_/ ∑_*j*_ 𝑝_$_ for 𝑗 ≠ 𝑖. In reality, the map 𝜑 cannot be perfectly learned, and the composition prediction always contains some error. To take this into account, we can compute the null composition 𝒑̅ by renormalizing the relative abundances of the remaining species from the predicted composition of the original community, i.e., 𝑝̅_*j*_ = 𝑝^…^_*j*_/ ∑_$%_ 𝑝^&^, where 𝒑^&^ = (𝑝^&^) = 𝜑(𝒛). This way, the prediction errors in 𝒑̃ and 𝒑̅ will be canceled to some extent, and hence the predicted keystoneness will be more accurate (see **Fig.S1**).

We can quantify the impact of species-𝑖 ’s removal on 𝑠 in two different ways: (1) *structural impact*, defined as the dissimilarity between the taxonomic profiles 𝒑̃ and 𝒑̅, i.e., 𝑑(𝒑̃, 𝒑̅); (2) *functional impact*, defined as the dissimilarity between the functional profiles ***f̃*** and ***f̅***, i.e., *d*(***f̃f̅***). Here ***f̃*** (or 𝒇̃) can be computed from 𝒑̃ (or 𝒑̅) and the genomic content network (GCN)^37^, respectively. The GCN is a weighted bipartite graph connecting the 𝑁 species to their genes (see **SI Sec.4,5**). Suppose there are in total 𝑀 genes in the metagenome of the 𝑁 species. The GCN can then be represented by an incidence matrix 𝑮 = (𝐺_*ia*_) ∈ ℝ^*N*×*M*^), where a non-negative integer 𝐺_*ia*_ indicates the copy number of gene-𝑎 in the genome of species-𝑖. The gene composition or functional profile 𝒇 ∈ Δ^)^ of a microbiome sample with taxonomic profile 𝒑 can then be calculated as 𝒇 = 𝑐 𝒑 𝑮, where *c* = [∑_*a=1*_^*M*^ ∑_*i*=1_^*N*^ *p_i_ G_ia_*]^−1^ is a normalization constant.

We define the *structural keystoneness* of species-𝑖 in a community 𝑠 = (𝒛, 𝒑) as

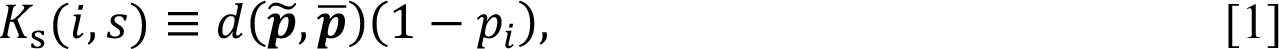

where 𝑑(𝒑̃, 𝒑̅) quantifies the structural impact of species-𝑖’s removal on the community 𝑠, while (1 − 𝑝_*i*_) captures how disproportionate is this effect. Note that 𝑑(𝒑̃, 𝒑̅) can be any distance or dissimilarity measure, e.g., the Bray-Curtis dissimilarity. 𝐾_-_(𝑖, 𝑠) = 0 indicates that the removal of species-𝑖 does not impact the abundance of any other species in the community 𝑠 at all. Similarly, we define the *functional keystoneness* of species-𝑖 in a community 𝑠 = (𝒛, 𝒑) as

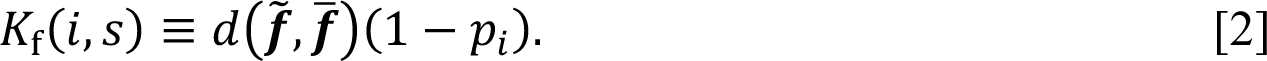

We emphasize that the structural (or functional) keystoneness defined here is community-specific, which is fundamentally different from those topological indices used in the food web and other ecological systems^38^.

### Validation of DKI framework

To demonstrate DKI’s performance in the keystoneness prediction, we generated synthetic data using the Generalized Lotka-Volterra (GLV) model with 𝑁 = 100 species in the species pool (meta-community). The initial species collection of each sample (local community) consisting of 50 species randomly drawn from the species pool (see **SI Sec.1**). We characterized the population dynamics of the meta-community using two parameters: (1) The connectivity 𝐶 of the underlying ecological network (which encodes all the pairwise inter-species interactions), representing the probability that two species interact directly. (2) The characteristic interaction strength 𝜎 represents the typical impact of one species over the per-capita growth rate of another species if they interact.

For a randomly generated ecological network with a random mixture of interaction types (e.g., parasitism, commensalism, mutualism, amensalism, or competition), without any heterogeneity in the network structure and links weights (i.e., interaction strengths), we expect that species keystoneness will not have any heterogeneity either. In other words, there will be no outlier or keystone species. To introduce keystone species to the local communities, inspired by a previous study^29^, we amplified all the link weights (inter-species interactions)^29^ to 𝑎̃_*ij*_ = 𝜃_*ij*_𝑎_*ij*_, where 𝜃_*ij*_ is randomly drawn from a log-normal distribution with mean 0 and standard deviation 𝜂. This will generate a few strong interactions, presumably leading to a few species with high keystoneness. Hereafter, we refer 𝜂 as the *characteristic amplification coefficient*.

We first trained cNODE2 to minimize the loss function defined as the mean Bray-Cutis dissimilarity between the true and predicted compositions for all samples (see **SI Sec.2**). Then, we evaluated DKI using all possible new species collection 𝒛̃ by removing each of present species in each sample. We systematically evaluated the performance of DKI in predicting the structural keystoneness 𝐾_*s*_ using simulated data generated from the GLV model with different values for the parameter pair (𝐶, 𝜂). We found that DKI can accurately predict the 𝐾_*s*_ of each species over different samples for a wide range of 𝐶 or 𝜂 values (see **Fig.2a-f**). The Spearman correlation 𝜌 between the true 𝐾_*s*_ (calculated from the simulated species removal process in the GLV model) and the predicted 𝐾_*s*_ is around 0.97 with p-value<0.001 (based on the AS 89 algorithm^39^).

**Figure 2:**
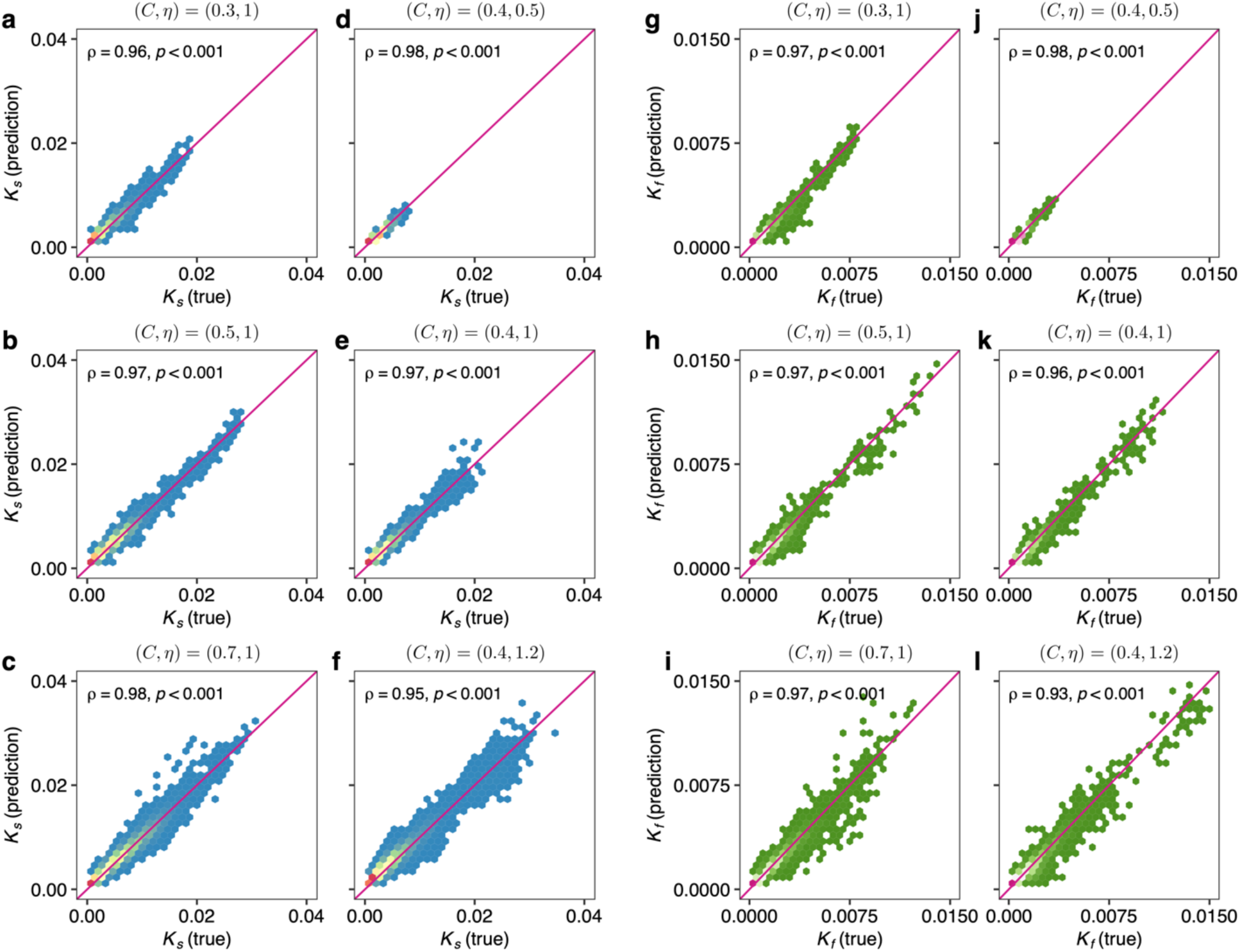
In silico validation of the DKI framework. Results are obtained for the pools of 𝑁 = 100 species with Generalized Lotka-Volterra population dynamics. We generate 500 samples to validate the performance of DKI. The population dynamics is characterized by two parameters: the network connectivity 𝐶 > 0 and boosting strength 𝜂 > 0. Predicted structural keystoneness and true structural keystoneness with network connectivity 𝐶 = 0.3 (a), 𝐶 = 0.5 (b) and 𝐶 = 0.7 (c) or boosting strength 𝜂 = 0.5 (d), 𝜂 = 1 (e) and 𝜂 = 1.2 (f). Predicted functional keystoneness and true functional keystoneness with network connectivity 𝐶 = 0.3 (g), 𝐶 = 0.5 (h) and 𝐶 = 0.7 (i) or boosting strength 𝜂 = 0.5 (j), 𝜂 = 1 (k) and 𝜂 = 1.2 (l). For different network connectivities, characteristic interaction strength 𝜎 = 0.01 and boosting strength 𝜂 = 1. For different boosting strengths, characteristic interaction strength 𝜎 = 0.01 and network connectivity 𝐶 = 0.4. In each panel, we show the Spearman correlation (𝜌) between the predicted and true keystoneness values, and the p-value.

To calculate the functional keystoneness using the synthetic data, we generated a random GCN displaying nested structure (with Nestedness metric based on Overlap and Decreasing Fill^40^ NODF = 0.31) for 100 species and 500 genes. We found that DKI can also accurately predict the 𝐾_0_of each species over different samples for a wide range of 𝐶 or 𝜂 values (see **Fig.2g-l**). The Spearman correlation 𝜌 between the true 𝐾_0_ (calculated from the simulated species removal process in the GLV model and the randomly generated GCN) and the predicted 𝐾_0_ is around 0.96 with p-value<0.001.

We emphasize that each species’ structural (or functional) keystoneness is context-dependent or community-specific. Yet, existing methods, especially those based on topological indices of correlation (or ecological) networks constructed (or inferred) from a collection of samples, cannot offer community-specific keystoneness. Moreover, those topological measures do not correlate with each species’ structural (or functional) keystoneness. To demonstrate this point, we generated synthetic data (see **SI Sec.1**) and compared the structural keystoneness 𝐾_-_(calculated from the simulated species removal process in the GLV model) with two classical topological indices, i.e., degree (the number of species connected with the species under consideration) and betweenness (the frequency of the species under consideration on the shortest paths connecting all pairs of other species) in the directed ecological network or the undirected correlation network. As shown in **Fig.3** **and** **Fig.4**, the two topological indices do not correlate with 𝐾_-_ at all, regardless of using the correlation network (**Fig.3**) or the ecological network (**Fig.4**).

**Figure 3:**
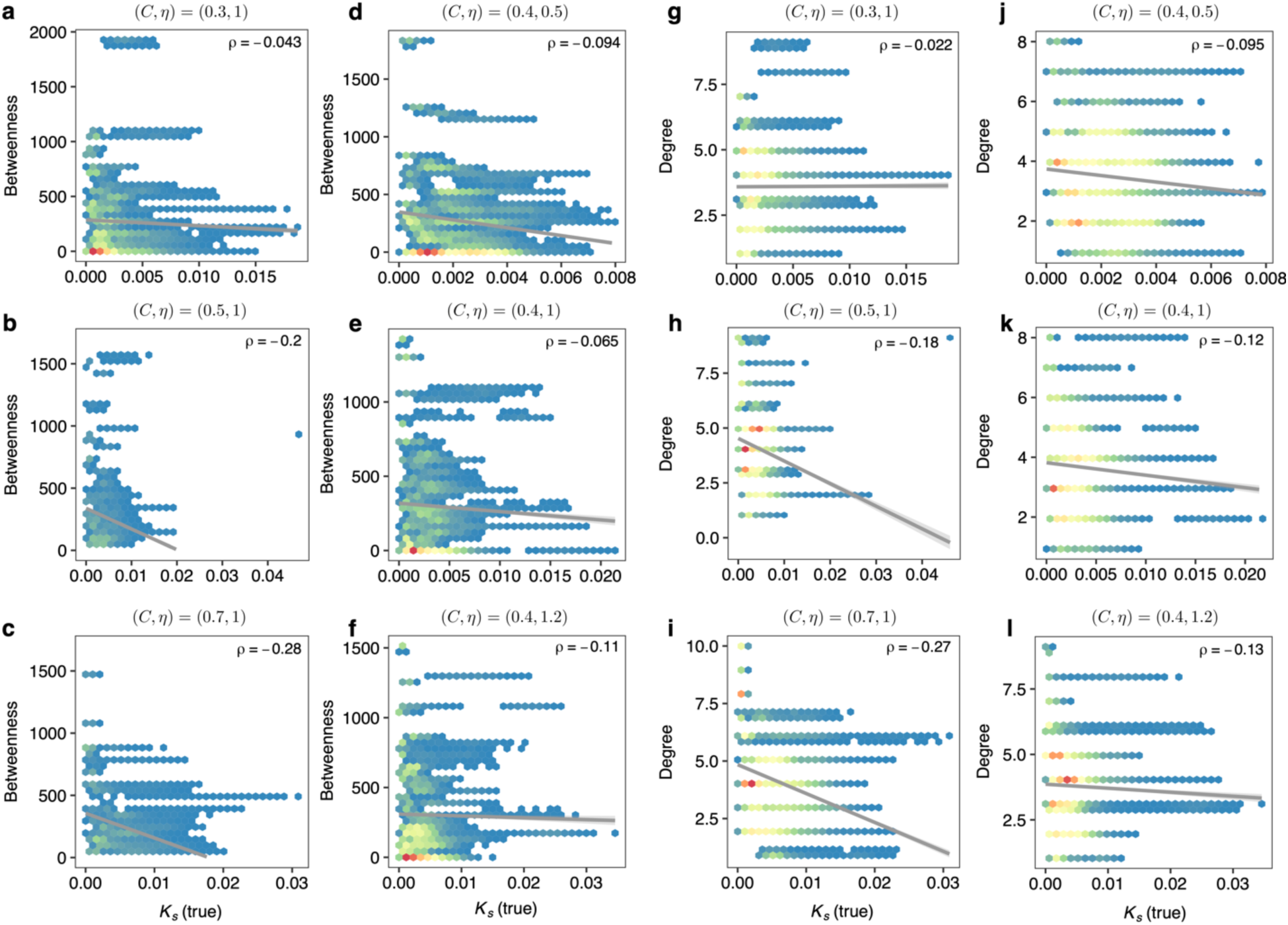
Traditional topological indices calculated from the undirected correlation network do not correlate with structural keystoneness. Synthetic samples (taxonomic profiles) were generated from the GLV model with 𝑁 = 100 species in the species pool. The initial species collection of each sample (local community) consists of 50 species randomly drawn from the species pool (see SI Sec.1). The structural keystoneness 𝐾_-_ of each species in each sample was calculated from the simulated species removal process in the GLV model. Two traditional topological indices (betweenness and degree) of each species were calculated from the correlation network of species abundances constructed using sparCC^16^ with threshold 0.1. a-c, Structural keystoneness vs. betweenness. The ecological network connectivity 𝐶 = 0.3 (a), 𝐶 = 0.5 (b), 𝐶 = 0.7 (c). The characteristic interaction strength 𝜎 = 0.01 and the boosting strength 𝜂 = 1. d-f, Structural keystoneness vs. betweenness with 𝜂 = 0.5 (d), 𝜂 = 1.0 (e) and 𝜂 = 1.2 (f). 𝐶 = 0.4 and 𝜎 = 0.01. g-i, Structural keystoneness vs. degree. 𝐶 = 0.3 (g), 𝐶 = 0.5 (h), 𝐶 = 0.7 (i). 𝜎 = 0.01 and 𝜂 = 1. j-l, Structural keystoneness vs. degree with 𝜂 = 0.5 (j), 𝜂 = 1.0 (k) and 𝜂 = 1.2 (l). 𝐶 = 0.4 and 𝜎 = 0.01.

**Figure 4:**
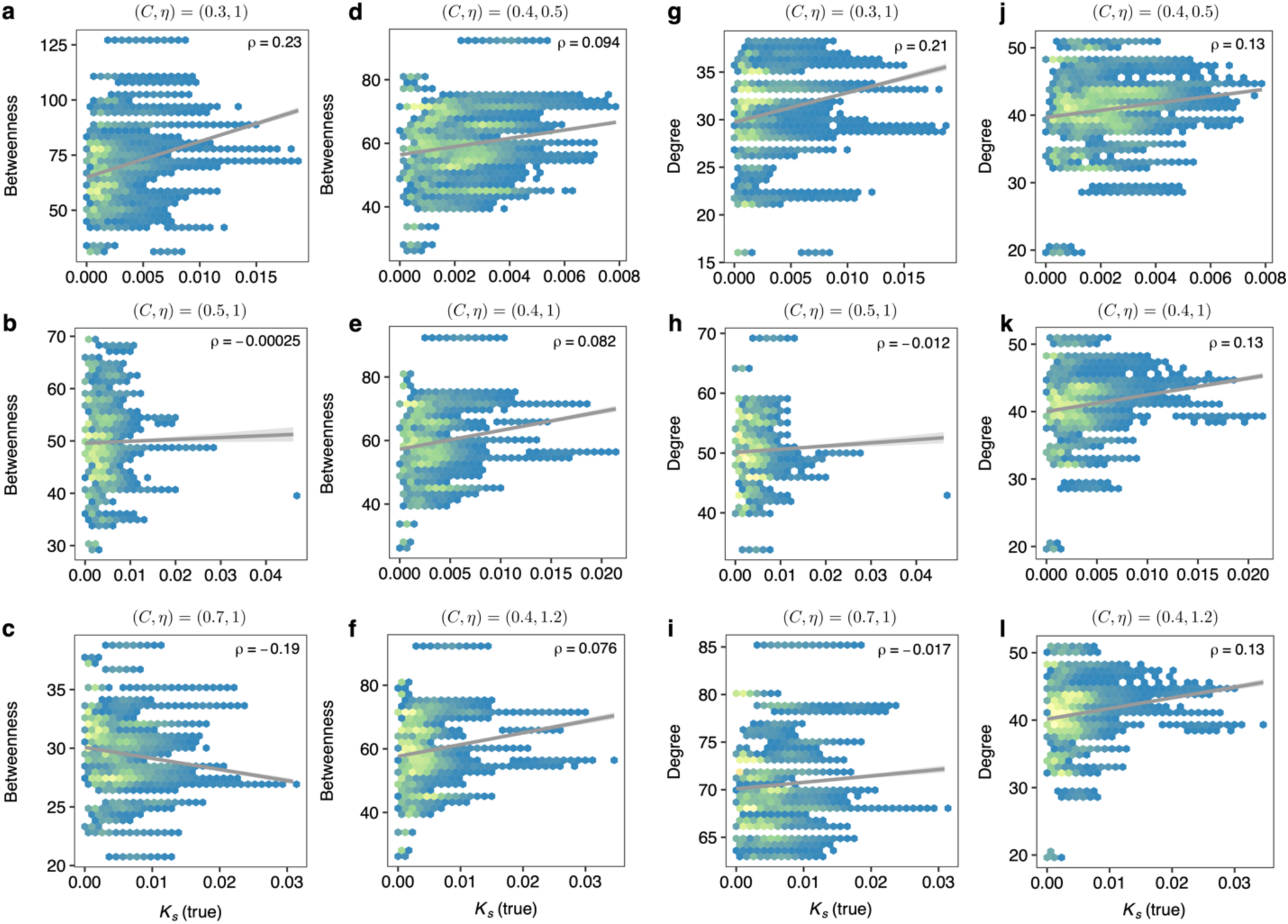
Traditional topological indices calculated from the directed ecological network do not correlate with structural keystoneness. Synthetic data and parameters in the GLV model are the same as used in the corresponding panels of Figure 3.

### Keystone species in the human gut microbiome

We applied the DKI framework to the human gut microbiome data collected in a curated metagenomic database^30^. We focused on the metagenomic data of stool samples of healthy adults aged between 18 to 65 and without antibiotics usage. In total, we have 2,815 samples involving 1,103 species.

We first trained cNODE2 by using all the 2,815 samples. Then for each of the 2,815 samples, we computed the structural keystoneness 𝐾_*s*_ for each species present in the sample. For species present in at least 10% of the samples, we ranked them based on their median structural keystoneness: median(𝐾_*s*_). **Fig.5a,b** show the top-20 and bottom 20 species, respectively. We found that those species with a higher median(𝐾_*s*_), e.g., *Prevotella copri*, tend to have a larger variation of their 𝐾_-_ across different samples, suggesting a stronger community specificity (**Fig.5a**); while those species with a lower median(𝐾_*s*_), e.g., *Alistipes finegoldii*, tend to have a smaller 𝐾_*s*_ variation, suggesting a weaker community specificity (**Fig.5b**).

**Figure 5:**
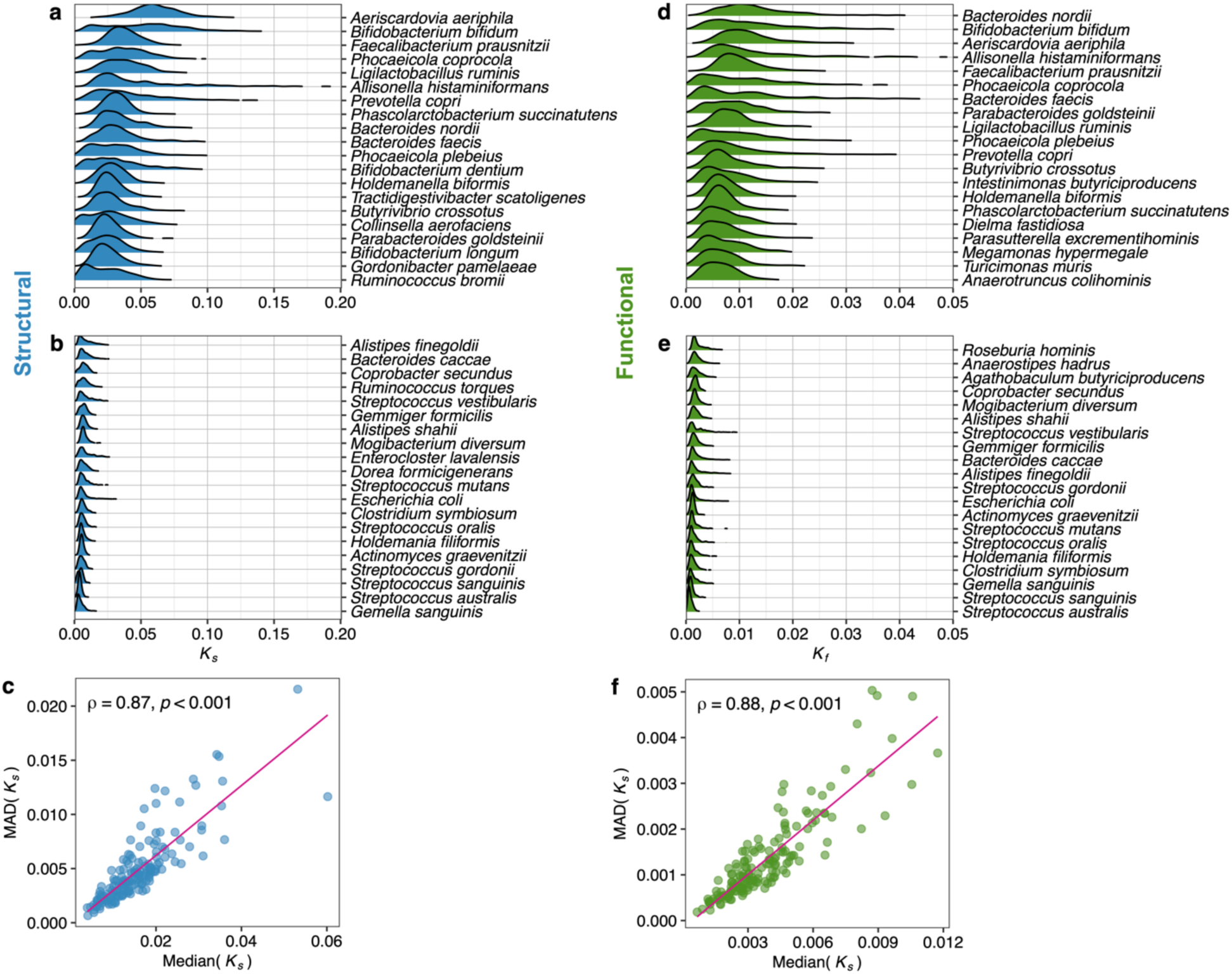
Keystone species in the human gut microbiome. We applied DKI to a large-scale human gut microbiome dataset collected in curatedMetagenomicData^30^. This dataset includes 2,815 fecal samples from healthy adults without antibiotics usage. In total, we have 1,103 species. **a-b,** Structural keystoneness distribution of the top-20 (a) and bottom-20 species (b) ranked by their median structural keystoneness. **c,** Spearman correlation between the median structural keystoneness and median deviation of keystoneness of each species. **d-e,** Functional keystoneness distribution of the top-20 (d) and bottom-20 species (e) ranked by their median functional keystoneness. **f,** Spearman correlation between the median functional keystoneness and median deviation of keystoneness of each species. In panels (a), (b), (d) and (e), the top/bottom-20 species were selected among species present in at least 10% of total samples.

To systematically explore the community-specificity of those species’ structural keystoneness, we plotted their median keystoneness median(𝐾_*s*_) vs. their median absolute deviation of structural keystoneness MAD(𝐾_*s*_) over all samples. We found that MAD(𝐾_*s*_) is highly correlated with median(𝐾_*s*_) with Spearman correlation 𝜌 = 0.87, p-value<0.001 (**Fig.5c**). This result indicates that taxa with low median structural keystoneness are unlikely to be keystone taxa in any community. By contrast, taxa with high median keystoneness have high keystoneness (and hence are likely keystone taxa) in some communities, but they can also have small keystoneness in other communities.

To compute the functional keystoneness, we constructed a reference GCN (See **SI. Sec.4** for details). We found that in general a species’ functional keystoneness is smaller than its structural keystoneness (see **Fig.S2**). This is closely related to the concept of functional redundancy^37^, i.e., multiple phylogenetically unrelated taxa can carry similar genes and perform similar functions. Similar to our results on structural keystoneness, we found that those species with a higher median(𝐾_0_), e.g., *Bifidobacterium bifidum*, tend to have a larger variation of their 𝐾*_f_* across different samples, suggesting a stronger community specificity (**Fig.5d**), while those species with a lower median(𝐾_0_), e.g., *Roseburia hominis*, tend to have a smaller 𝐾_0_ variation, suggesting a weaker community specificity (**Fig.5e**). We found that median(𝐾_0_) and MADH𝐾_0_I are strongly correlated: Spearman correlation 𝜌 = 0.88, p-value<0.001 (**Fig.5f**).

Based on the ranking of the median structural keystoneness, we found that among those top-ranking species, many of them have been identified as keystone species that carry unique functions and are essential for the maintaining host-microbe hemeostasis^41^. For example, Bifidobacteria are keystone microorganisms in gut microbiota associated with early life^42^; *Prevotella copri* is a keystone species of a healthy human intestinal mucosa^43^; *Faecalibacterium prausnitzii* is a keystone species that produces butyrate and its reduced abundance has been associated with Crohn’s disease^44^; *Bifidobacterium longum* is a minority species, but influences gut microbiota formation by breaking down complex carbohydrates and providing degradants for other bacterial groups to use^45^; *Ruminococcus bromii* plays key roles in promoting the synergistic utilization of resistant starch (RS) by initiating degradation of insoluble RS particles^5, 41^ (**Fig.5a**).

Interestingly, some of the species with high median structural keystoneness also have high median functional keystoneness, e.g., *Faecalibacterium prausnitzii* and *Prevotella copri* (**Fig.5d**). Based on the high median functional keystoneness, we also identified some potential keystone species that have been reported to perform important functions. For example, *Intestinimonas*-like bacteria are important butyrate producers that utilize N-ε-fructosyllysine and lysine in formula-fed infants and adults^46^.

### Keystone species in the human oral microbiome

We applied the DKI framework to the data collected in a large-scale human oral microbiome study^31, 47, 48^. We focused on the metagenomic data of saliva samples of healthy adults without any dental disease in the 4D-SZ cohort of this study^31, 47, 48^. In total, we have 1,412 samples involving 702 species.

We first trained cNODE2 by using all the 1,412 samples. Then for each of the 1,412 samples, we computed the structural keystoneness 𝐾_*s*_for each species present in the sample. To compute the functional keystoneness, we performed the genome-wide functional annotation using eggNOG mapper^49^, and then constructed GCN by calculating the copy number of gene products for each species.

For species present in at least 10% of the samples, we ranked them based on their median structural (or functional) keystoneness: median(𝐾_*s*_) (or median(𝐾_0_), respectively). **Fig.6a,b** (and **Fig.6d,e**) show the top 20 and bottom 20 species based on their median(𝐾_*s*_) (or median(𝐾_0_)), respectively. We found that the community-specificity of both structural and functional keystoneness (quantified by MAD(𝐾_*s*_) and MADH𝐾_0_I, respectively) in the oral microbiome is weaker than that observed from the gut microbiome. But the median structural (or functional) keystoneness is still correlated with the median absolute deviation of structural (or functional) keystoneness over all samples: 𝜌 = 0.67 (or 0.63) for structural (or functional) keystoneness, p-value<0.001 in both cases (**Fig.6c,f**).

**Figure 6:**
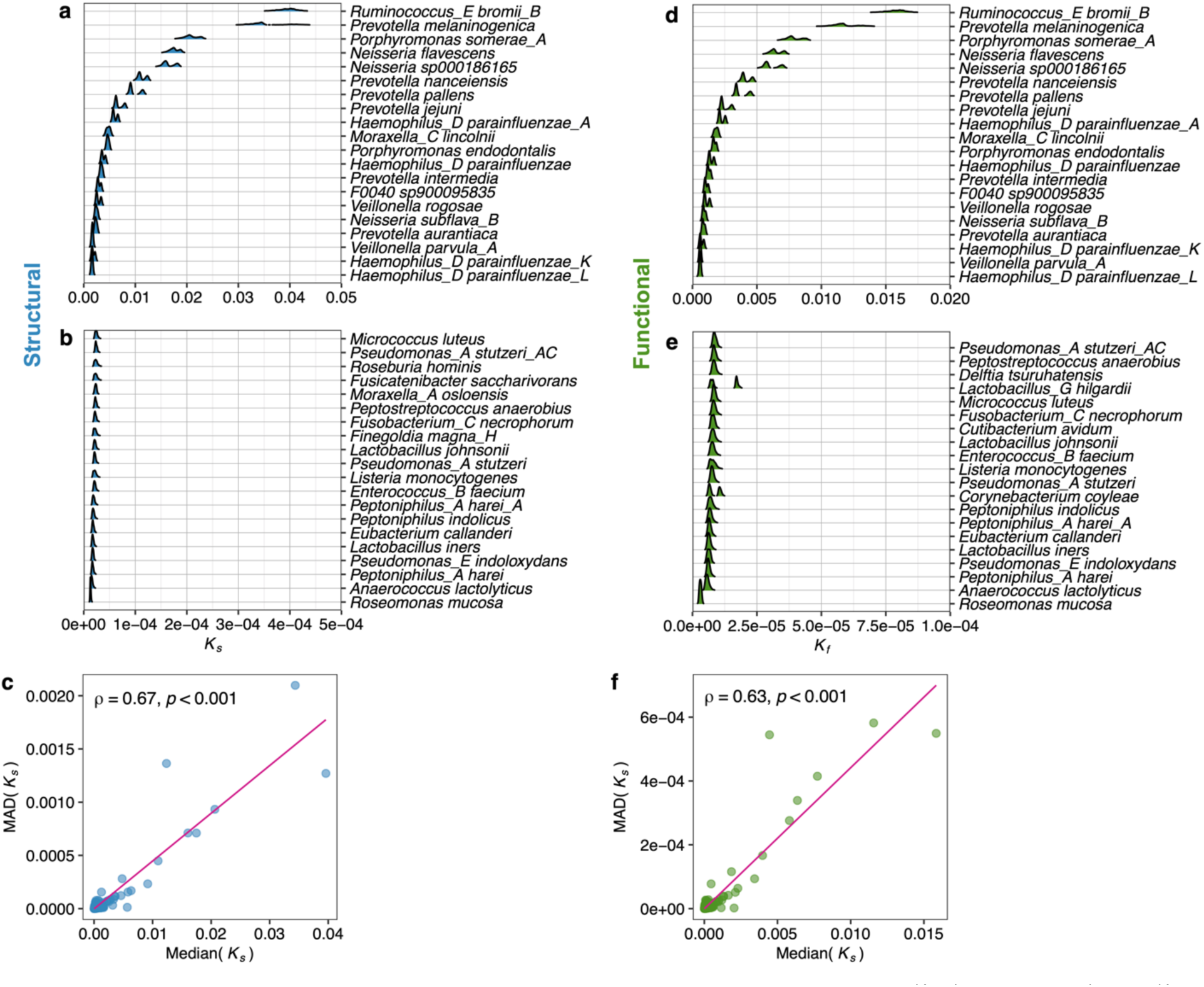
Keystone species in the human oral microbiome. We applied DKI to the saliva microbiome data of healthy adults in a large-scale human oral microbiome study^31^. The dataset includes 1,421 saliva samples with a total of 702 species. **a-b,** Structural keystoneness distribution of the top-20 (a) and bottom-20 species (b) ranked by their median structural keystoneness. **c,** Spearman correlation between the median structural keystoneness and median deviation of keystoneness of each species. **d-e,** Functional keystoneness distribution of the top-20 (d) and bottom-20 species (e) ranked by their median functional keystoneness. **f,** Spearman correlation between the median functional keystoneness and median deviation of keystoneness of each species. In panels (a), (b), (d) and (e), the top/bottom-20 species were selected among species present in at least 10% of total samples.

Among those top-ranking species, some species are members of the genus *Prevotella*, which is one of the core anaerobic genera in the oral microbiome involved in gastrointestinal and respiratory health and disease^50^. Two species are members of the genus *Veillonella*, which plays an important role in oral biofilm ecology, e.g., establishing mutualistic relationships with other members of the oral microbiome to contribute to the health equilibrium^51^. The genus *Neisseria* is associated with oral disease-free individuals, and an increase in *Neisseria* can be considered a positive change in the microbiota related to general oral health^52^.

### Keystone taxa in the environmental microbiomes

Finally, we applied the DKI framework to two large-scale environmental microbiome datasets: (1) A soil microbiome dataset^32^ with 1,160 soil microbiome samples collected from Central Park in New York City^32^. To ensure the sample size is comparable to the number of taxa, we focused on the analysis of taxonomic profiles at the order level (with 298 orders in total). (2) A coral microbiome dataset^33^ with 1,400 coral microbiome samples. We focused on the analysis of genus-level taxonomic profiles (with 1,054 genera in total).

Following the same procedure as above, we first trained cNODE2 by using all the 1,160 soil (or 1,400 coral) microbiome samples, respectively. Then for each of the soil (or coral) microbiome samples, we computed the structural keystoneness 𝐾_*s*_for each order (or genus) present in the sample, respectively. For those taxa that are present in at least 10% of the samples, we ranked them based on their median structural keystoneness: median(𝐾_*s*_). Similar to we observed from the human gut microbiome data analysis, the structural keystoneness is strongly (or weakly) context-dependent for taxa with higher (or lower) median keystoneness for both soil (**Fig.7a-b**) and coral (**Fig.8a-b**) microbiome; and MAD(𝐾_*s*_) is highly correlated with median(𝐾_*s*_)with Spearman correlation 𝜌 = 0.79, p-value<0.001 for soil microbiome (**Fig.7c**) and 𝜌 = 0.93, p-value<0.001 for coral microbiome (**Fig.8c**).

To evaluate functional keystoneness, we used PICRUSt2^53^ to predict the KO content of each ASV (see SI Sec.5 for details). Then the KO content of each order (or genus) was obtained by averaging the KO contents of all ASVs annotated to that order (or genus) for the soil (or coral) microbiome, respectively. Again, we found the functional keystoneness is strongly (or weakly) context-dependent for taxa with higher (or lower) median keystoneness for both soil (**Fig.7:d-e**) and coral (**Fig.8:d-e**) microbiome. Moreover, the median(𝐾_0_) is positively correlated with MAD(𝐾_0_) with Spearman correlation 𝜌 = 0.95, p-value<0.001 for both soil (**Fig.7f**) and coral (**Fig.8f**) microbiome.

**Figure 7:**
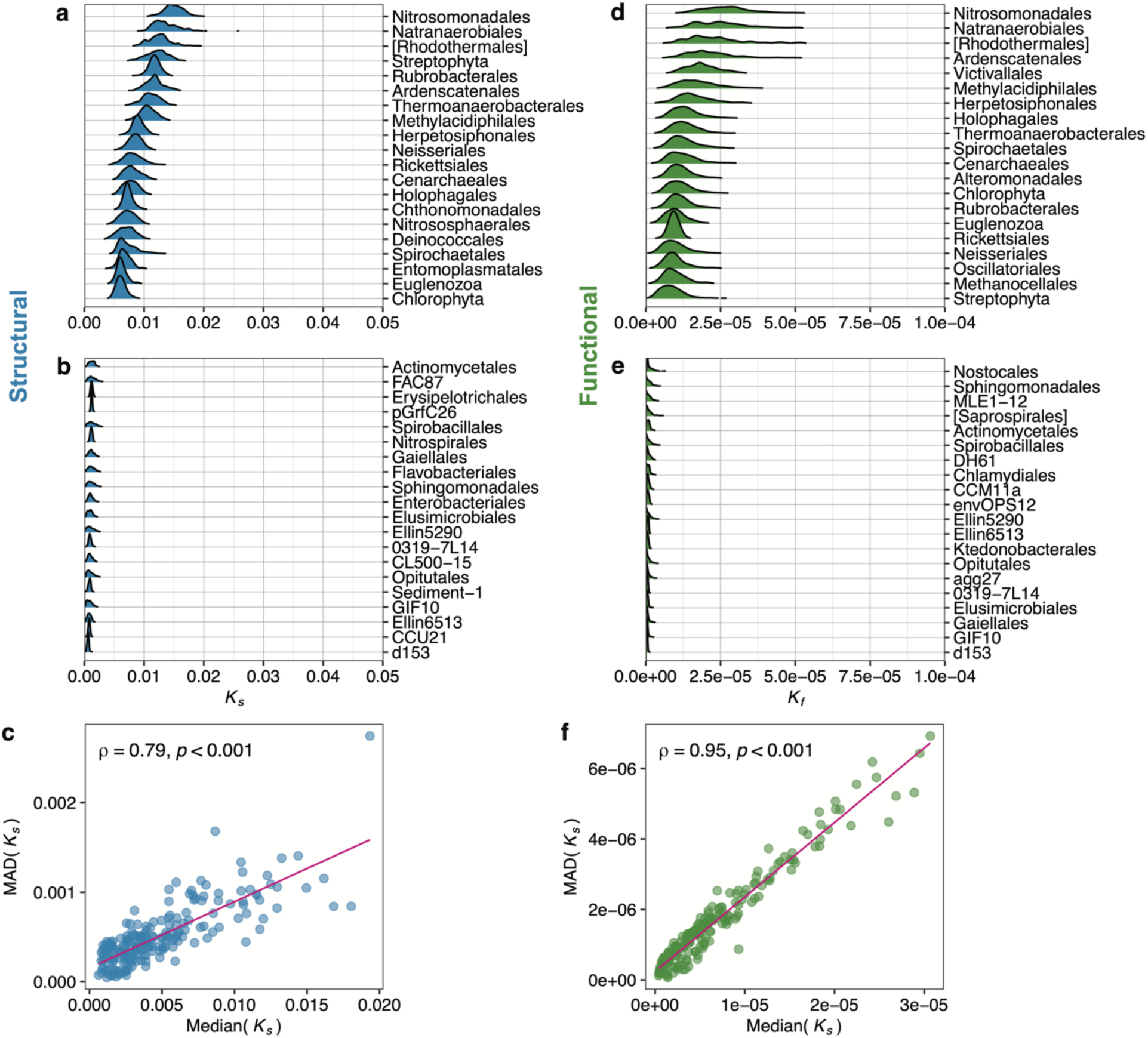
Keystone orders in the soil microbiome. We applied DKI to a large-scale soil microbiome dataset^32^. The dataset includes 1,160 soil samples with a total of 298 orders. **a-b,** Structural keystoneness distribution of the top-20 (a) and bottom-20 orders (b) ranked by their median structural keystoneness. **c,** Spearman correlation between the median structural keystoneness and median deviation of keystoneness of each order. **d-e,** Functional keystoneness distribution of the top-20 (d) and bottom-20 orders (e) ranked by their median functional keystoneness. **f,** Spearman correlation between the median functional keystoneness and median deviation of keystoneness of each order. In panels (a), (b), (d) and (e), those top/bottom-20 orders were selected among orders present in at least 10% of total samples.

**Figure 8:**
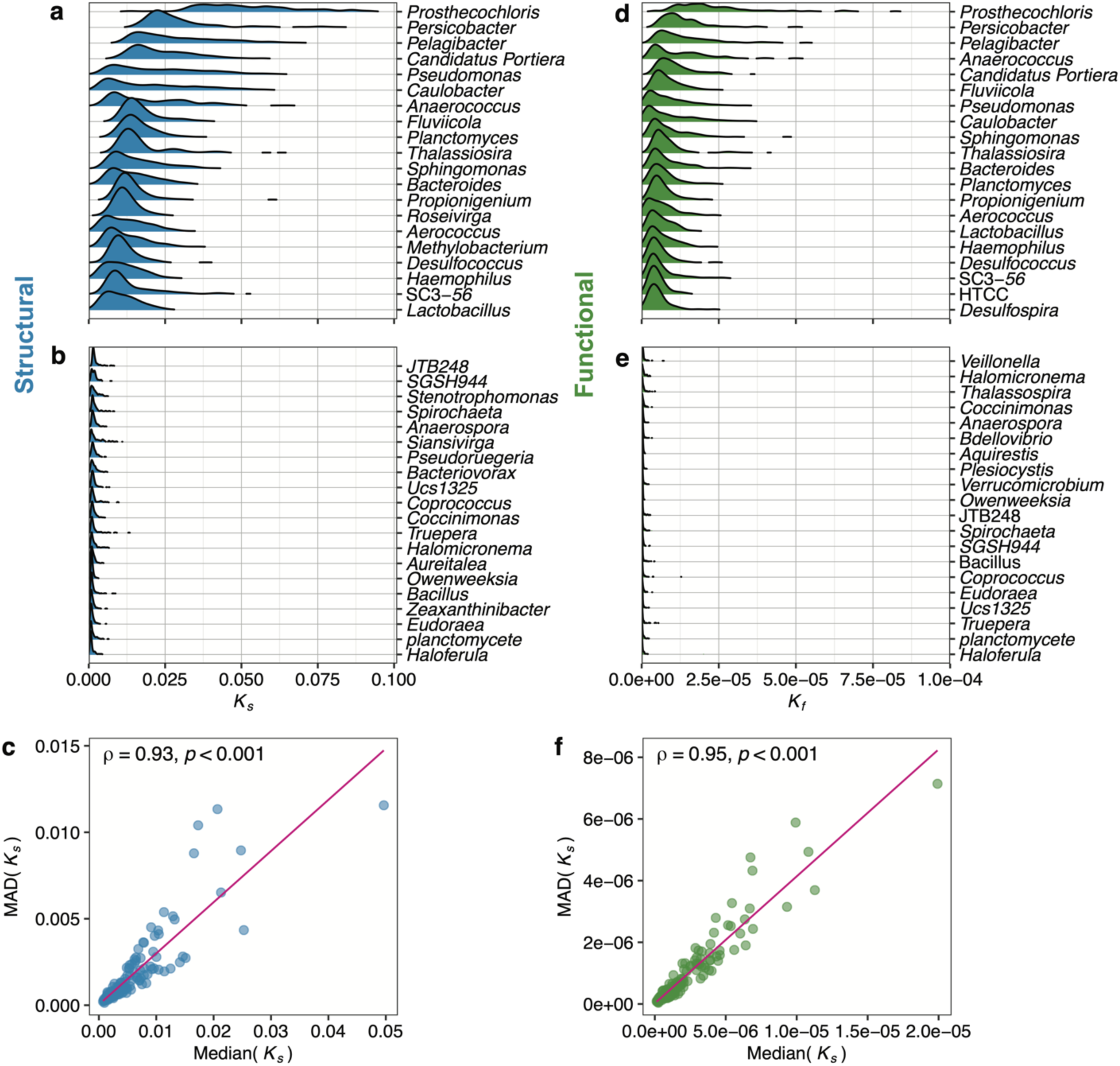
Keystone genera in the coral microbiome. We applied DKI to a large-scale coral microbiome dataset^33^. The dataset includes 1,400 samples with a total of 1,054 genera. **a-b,** Structural keystoneness distribution of the top-20 (a) and bottom-20 genera (b) ranked by their median structural keystoneness. **c,** Spearman correlation between the median structural keystoneness and median deviation of keystoneness of each genus. **d-e,** Functional keystoneness distribution of the top-20 (d) and bottom-20 genera (e) ranked by their median functional keystoneness. **f,** Spearman correlation between the median functional keystoneness and median deviation of keystoneness of each genus. In panels (a), (b), (d) and (e), the top/bottom-20 genera were selected among genera present in at least 10% of total samples.

For the soil microbiome, among the top-ranking orders (based on structural or functional keystoneness), we found several potential keystone taxa. For example, Nitrosomonadales is an order critical to the biogeochemical cycling of nitrogen, sulfur, carbon, and iron. Many species in this order play key roles in principal biochemical processes^54^. Alteromonadales is a putative keystone order associated with hydrocarbon reservoirs^55^. Thermoanaerobacterales is the most affected order that appeared in the soil microbial community after the addition of crop residue, which can drive the soil organic carbon and nitrogen dynamics in the presence of the crop residues^56^.

For the coral microbiome, among those top-ranking genera (based on structural and/or functional keystoneness), *Prosthecochloris* is a genus of green sulfur bacteria living in coral reefs, which are crucial in generating available nitrogen in the nutrient-limited environment^57^. The genus *Pseudomonas* has been demonstrated to play a central role in the degradation of dimethylsulfoniopropionate (DMSP) to dimethyl sulfide (DMS) and acrylic acid^58, 59^.

## Discussion

The concept of keystone species has been extensively investigated in ecology. Despite the considerable confusion and difference^3^, the operational definitions agree that a keystone species disproportionately affects its natural environment relative to its abundance. Systematically identifying keystone species in complex microbial communities is very challenging due to our limited knowledge of the population dynamics of those communities, as well as many logistical and ethical concerns regarding the manipulation of those communities. In this work, we propose a data-driven framework to systematically identify keystone species in complex microbial communities. This framework enables us to compute the structural and functional keystoneness of each species in a community for the first time. Our framework can be used to facilitate data-driven management of complex microbial communities.

We emphasize that the proposed framework is general enough and can be modified in many different ways. For example, instead of using cNODE2, one can use other more advanced deep learning models to learn the map 𝜑: 𝒛 ↦ 𝒑, upon their availability. One can use other dissimilarity measures to quantify the structural or functional impact of species’ removal, instead of using Bray-Curtis dissimilarity. One can also use other formulas that combine the impact component and the biomass component to quantify the keystoneness^60^. In addition, beyond studying the impact of a species’ removal on the community-level functional profile, we can also focus on its impact on any specific microbial function and hence quantify its vulnerability. For instance, one can calculate the relative abundances of a function before and after a species’ removal, respectively.

There are several caveats in applying the proposed DKI framework. We know that the performance of DKI is largely determined by the accuracy of learning the map 𝜑: 𝒛 ↦ 𝒑. To achieve high accuracy in learning the map, we typically need a large set of microbiome samples to adequately train the deep learning model. Based on our previous experience^36^, for a metacommunity of 𝑁 taxa, about 2𝑁 samples are required. In addition, the dataset should meet the following assumptions: (1) the metacommunity has strongly universal microbial dynamics; (2) the compositions of the microbiome samples represent (at least approximately) the steady states of the microbial communities; and (3) there is a unique steady-state composition associated with each species collection. Those assumptions are necessary to ensure the learning of the map 𝜑: 𝒛 ↦ 𝒑 is a mathematically well-defined problem.

## Acknowledgements

**Acknowledgments.** Y.-Y.L. acknowledges the funding support from National Institutes of Health (R01AI141529, R01HD093761, RF1AG067744, UH3OD023268, U19AI095219, and U01HL089856).

## Ethics declarations

**Competing financial interests.** The authors declare no competing financial interests.

Author Contributions. Y.Y.L. conceived and designed the project. X.W.W. performed all the numerical calculations. X.W.W. and Z.S. analyzed real data. X.W.W. and Y.Y.L. wrote the manuscript. All authors analyzed the results, edited, and approved the manuscript.

**Data accessibility.** The data and code used in this work are available at https://github.com/spxuw/DKI.

**Correspondence.** Correspondence and requests for materials should be addressed to Y.-Y. L.

**Identifying keystone species in microbial communities using deep learning**

## Supplementary Information

### Table of Contents

***1. Using an ecological model to generate synthetic microbiome data. 2***

***2. Prediction species composition from species assemblage 2***

***3. Existing definitions of keystoneness. 4***

3.1 Euclidean distance-based definition. 4

3.2 Topology and abundance-based definition 4

***4. Human microbiome datasets 4***

***5. Environmental microbiome datasets 6***

## **1.** Using an ecological model to generate synthetic microbiome data

To systematically examine the performance of cNODE in the prediction of keystoneness, we generated synthetic data using the classical generalized Lotka–Volterra (GLV) model^1^:

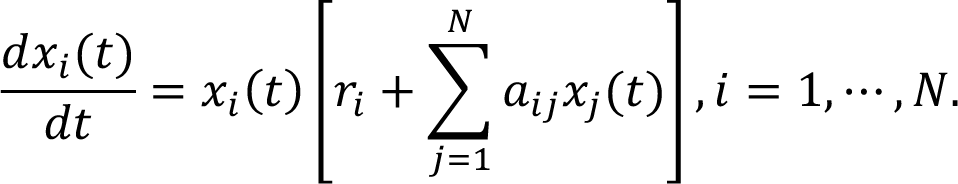

Here 𝑥_*i*_(𝑡) represents the absolute abundance of the *i*-th species at time 𝑡 ≥ 0. The pair-wise microbial interaction is presented by the matrix 𝐴 = (𝑎_*ij*_) ∈ ℝ^*N*×*N*^, with 𝑎_*ij*_ > 0 (< 0, or = 0) means that species-*j* promotes (inhibits or does not affect) the growth of species-*i*, respectively. The ecological network 𝐺(𝐴) is constructed using an Erdős-Rényi random graph model with 𝑁 nodes (species) and connectivity 𝐶 (the probability connecting two species). To generate the interaction matrix 𝐴 of ecological network, for each link (𝑗 → 𝑖) ∈ 𝐺(𝐴) with 𝑗 ≠ 𝑖, we draw 𝑎_*ij*_from the normal distribution ℕ(0, 𝜎). All other entries of 𝐴 are set to be zero. The intrinsic growth rate vector 𝑟 is drawn from uniform distribution 𝒰(0,1) for all species. The species collection of each local community (sample) was randomly drawn 50 species from the total 𝑁 = 100 pool. The initial state of each sample was randomly drawn from the uniform distribution 𝒰(0,1) and the final state was obtained after integrating the dynamics into a steady state.

To strengthen the heterogeneity of interactions among species, and hence “plant” keystone species to the ecological network, we introduced another interaction strength so that the interaction strength 𝑎_*ij*_between two species (𝑖, 𝑗) was amplified to 𝑎C_*ij*_ = 𝜃_*ij*_𝑎_*ij*_if they interact^2^ and 𝜃 is drawn from a lognormal distribution with mean 0 and standard deviation 𝜂. This will generate a few highly influential inter-species interactions whilst most other interactions have low interaction strength.

## **2.** Prediction species composition from species assemblage

The key idea of our Data-driven Keystone species Identification (DKI) framework is using a deep learning method to predict microbial compositions from species assemblages. Given a species pool Ω = {1, ⋯, 𝑁} associated with a particular habitat (e.g., the human gut), any microbiome sample collected from this habitat can be considered as a local community assembled from the species pool Ω. Consider a set of local communities collected from this habitat, e.g., gut microbiome samples from different hosts. The species assemblage of a sample can be represented by a binary vector 𝒛 ∈ {0,1}^*N*^, where its 𝑖-th entry satisfies 𝑧_*i*_ = 1 (𝑧_*i*_ = 0) if species 𝑖 is present (or absent) in this sample. The microbial composition of this sample is characterized by a vector 𝒑 ∈ Δ^*N*^, where its 𝑖-th entry 𝑝_*i*_ is the relative abundance of species 𝑖 in this sample and Δ^*N*^ is the probability simplex. Any deep learning methods, including Multi-Layer Perceptron^3^ (MLP) or ResNet^4^ can be used to learn such a map. In a previous study, we developed a deep learning architecture: cNODE^5^ (composition Neural Ordinary Differential Equation) to learn the map 𝜑: 𝒛 ∈ {0,1}^*N*^ ↦ 𝒑 ∈ Δ^*N*^ directly from a set of microbiome samples from a habitat without using any population dynamic model. As long as we have collected enough samples (to train the deep learning model cNODE), cNODE can predict the composition of any unseen species assemblage based on 𝒑 = 𝜑(𝒛).

To learn such a map, we first transform the species collection 𝒛 ∈ {0,1}^*N*^ into the initial condition 𝒉(0) = 𝒛/𝟏^T^𝒛 ∈ Δ^*N*^, where 𝟏 = (1, ⋯,1)^𝐓^, then the initial condition is used to solve the set of nonlinear ODEs:

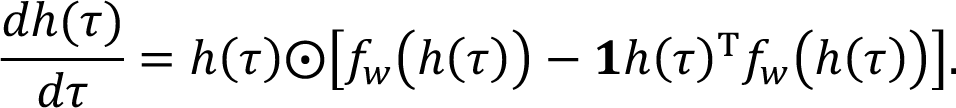

Here, 𝜏 ≥ 0 represents virtual time and ⨀ is the entry-wise multiplication. The function 𝑓_)_: Δ^*N*^→ ℝ^*N*^ is continuous function parameterized by 𝑤. Note that in the original cNODE architecture, we used a single layer linear function 𝑓_)_(ℎ) = 𝑤_%_ℎ to reduce the parameter space due to small sample size. However, the single-layer architecture is too simple to capture the complex, i.e., non-linear interactions between species. In the current study, we used a two-layer linear function 𝑓_)_(ℎ) = 𝑤_*_(𝑤_%_ℎ) and 𝑤_%_ ∈ ℝ^*N*×*N*^, 𝑤_*_ ∈ ℝ^*N*×*N*^ to learn more complex patterns in the data (see Figure 1). To avoid confusion, we called this deep learning architecture cNODE2. Those ODEs can be numerically integrated to obtain the prediction 𝒑^ = ℎ(𝜏_*c*_) and we choose 𝜏_+_ = 100. Thus, cNODE2 is used to learn the map

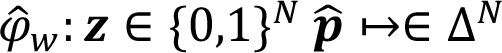

through species collection 𝒛 to predict the composition 𝒑^. cNODE2 is trained to minimize the loss function:

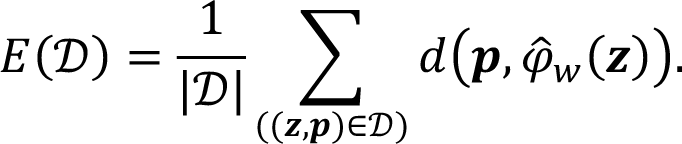

Here 𝒟 is the data set, and 𝑑 can be any distance or dissimilarity measure, e.g., the Bray-Curtis dissimilarity. We used different batch sizes for different datasets due to the sample size. The batch size for synthetic (or real) datasets is 20 (or 50), respectively. The total epoch is 1,000 for all datasets.

## **3.** Existing definitions of keystoneness

### 3.1 Euclidean distance-based definition

In the MDSINE (Microbial Dynamics Systems INference Engine) framework^6^, the keystoneness of a species is defined as follows. The measure starts from the community composition that allows the largest number of species to stably coexist, then to quantify the keystoneness of species-𝑖, we remove it from the community and calculate the steady-state concentrations after removing species-𝑖. The keystoneness of species-𝑖 is calculated as the Euclidean distance between the concentrations of the staring profiles and that of the profile with species-𝑖 removed, with the calculation excluding the contribution of the species-𝑖. Yet, this definition does not consider the disproportionate effort of a species’ keystoneness in terms of its abundance. In addition, most of microbiome datasets are compositional, thus the compositionality needs to be taken into account in the keystoneness calculation.

### 3.2 Topology and abundance-based definition

In the LIMITS (Learning Interactions from Microbial Time Series) framework^7^, keystone species are identified as follows. First, LIMITS is used to obtain a reliable estimate for the topology of the ecological network by employing spare linear regression with bootstrap. Then, the keystone species are those low-abundance species that exert tremendous influence on the structure of the microbial communities.

## **4.** Human microbiome datasets

To identify keystone species in the human gut microbiome, we leveraged the curatedMetagenomicData database^8^. Note that curatedMetagenomicData provides standardized, curated human microbiome data for downstream analyses. In particular, the taxonomic and functional profiles of all the microbiome samples in this database were calculated with MetaPhlAn2^9^ and HUMAnN2^10^ respectively. We focused on stool samples of healthy adults based on the following criteria: (1) age is between 18 and 65; (2) without antibiotic usage and (3) sequencing platform IlluminaHiSeq. This results in 2,815 samples involving 1,103 species. To construct the GCN and compute the functional keystoneness, we first generated the sample-specific GCN for each sample. The profiling results provided by curatedMetagenomicData include the contributions of various species to each gene abundance (generated by HUMAnN2^10^), rendering the construction of sample-specific GCNs relatively easy. In particular, for a given sample, we calculated the copy number of gene-𝑎 in the genome of species-𝑖, denoted as 𝐺_*ia*_, as the abundance of gene-𝑎 divided by the abundance of species-𝑖 in the sample. Then we merged all the sample-specific GCNs and use the average gene copy number 〈𝐺_*ia*_〉 in the final GCN. Note that when calculating the average copy number for a specific gene in the genome of a species, we only considered those samples with the corresponding species detected.

Unfortunately, the sample size of oral microbiome in curatedMetagenomicData is not big enough to train cNODE2. To identify keystone species in the human oral microbiome, we analyzed data collected in a large-scale human oral microbiome study^11^. This study used metagenomic shotgun data for 3,346 oral metagenomic samples together with 808 published samples to obtain 56,213 medium-and high-quality metagenome-assembled genomes (MAGs). The 56,213 new MAGs and 190,309 existing genomes were clustered into 3,589 species-level genome bins (SGBs). Taxonomic classification of the 3,589 SGBs was assigned using the GTDB-Tk (v0.3.2) classification workflow^12^ with external GTDB release 89. Genome-wide functional annotation was conducted by using EggNOG mapper (v1.0.3) through orthology assignment. For our analysis, we focused on the metagenomic data of saliva samples of healthy adults without any dental disease from the 4D-SZ cohort of this study. We initially filtered out those SGBs that did not have species-level annotations. Then, we gathered the species names of the remaining SGBs. In total, we have 1,412 samples involving 702 species. Since the genome information (NCBI assembly accession number) of their species is available through GTDB-TK annotation for the oral samples, the GCN here is thereby constructed through parsing the genome annotation file (e.g., the fnn files). Specifically, we counted the copy numbers of gene products annotated in the genome (excluding hypothetical proteins) to finally construct the GCN.

Because the taxonomic (and functional) profiles in the two studies^8, 11^ were calculated with different procedures, we did not use exactly the same method to construct the GCN associated with the human gut and oral microbiome. But we emphasize that the underlying strategy is conceptually the same.

## **5.** Environmental microbiome datasets

For the soil microbiome, we downloaded the taxonomic profiles from Qiita^14^ (ID: 2104). This dataset includes the 1,160 microbial communities of Central Park in New York City^14^. We summarized the ASV table into order levels. Taxa not annotated into order-level were merged and considered as an unsigned taxon. For the coral microbiome, we downloaded the taxonomic profiles from the Qiita^14^ (ID: 10895). This dataset includes 1,400 coral samples^15^. We summarized the ASV table into the genus-level and taxa not annotated into the genus-level were merged and considered as an unsigned taxon.

To compute the functional profiles, we used the entire PICRUSt2 pipeline (“picurst2_pipeline.py”) to generate the metagenome prediction for 16S data. The parameters used are all default values in PICRUSt2. We constructed the GCN using the predicted copy number of KO per ASV. The KO content of each order (or genus) was obtained by averaging the KO contents of all ASVs annotated to that order (or genus).

---

**Figure S1:**
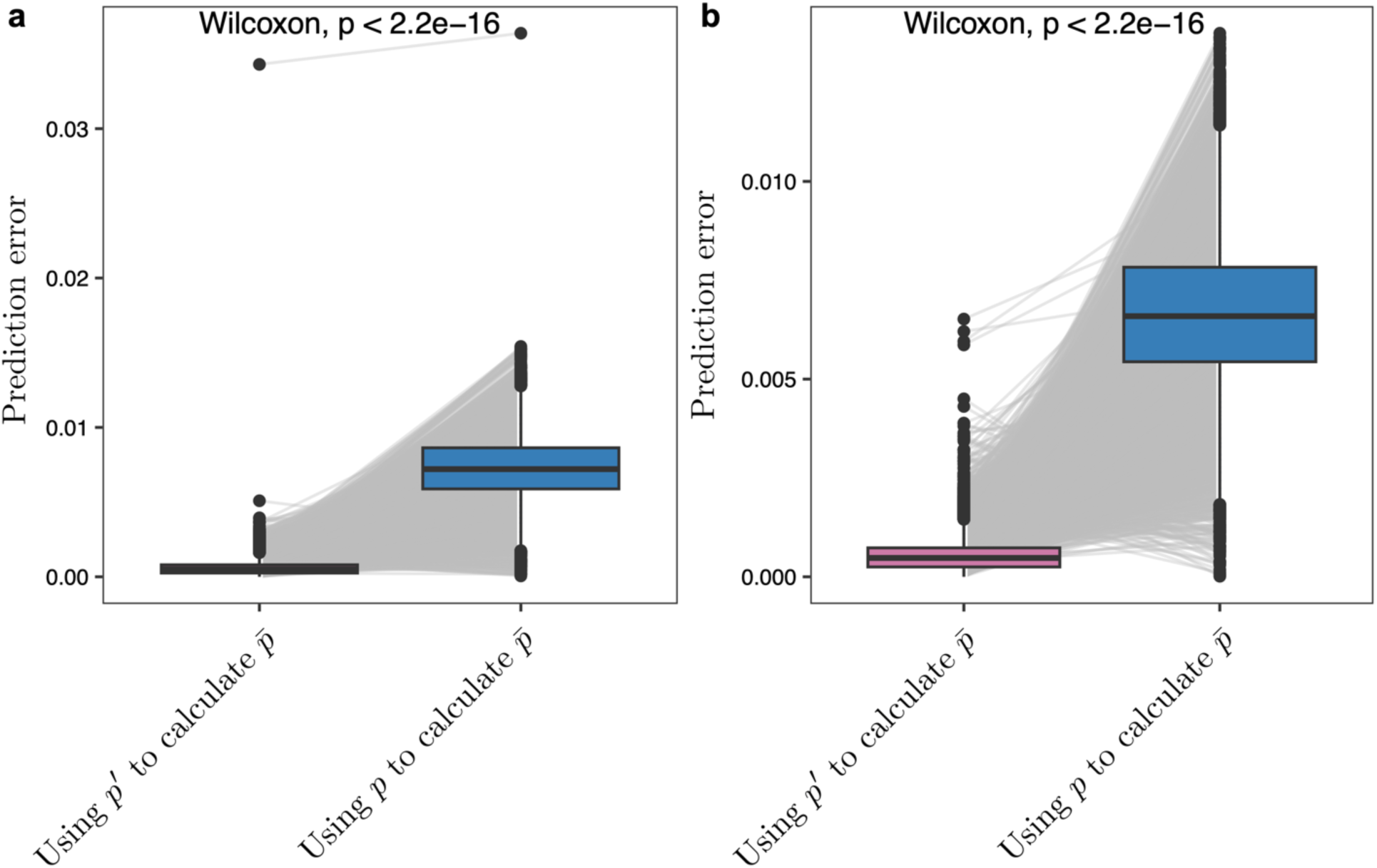
K**e**ystoneness **calculated from the predicted and true compositions.** We compared the keystoneness error defined as the L1 distance between the true keystoneness (calculated by running the GLV population dynamics model) and the predicted keystoneness using the true composition 𝑝 (blue) or the predicted composition 𝑝′ (pink), respectively. **a,** network connectivity 𝐶 = 0.5, characteristic interaction strength 𝜎 = 0.01 and boosting strength 𝜂 = 1. **b,** 𝜂 = 1, 𝜎 = 0.01 and 𝐶 = 0.4. Boxes indicate the interquartile range between the first and third quartiles with the central mark inside each box indicating the median. Whiskers extend to the lowest and highest values within 1.5 times the interquartile range. P-value was calculated using the paired Wilcoxon test.

**Figure S2:**
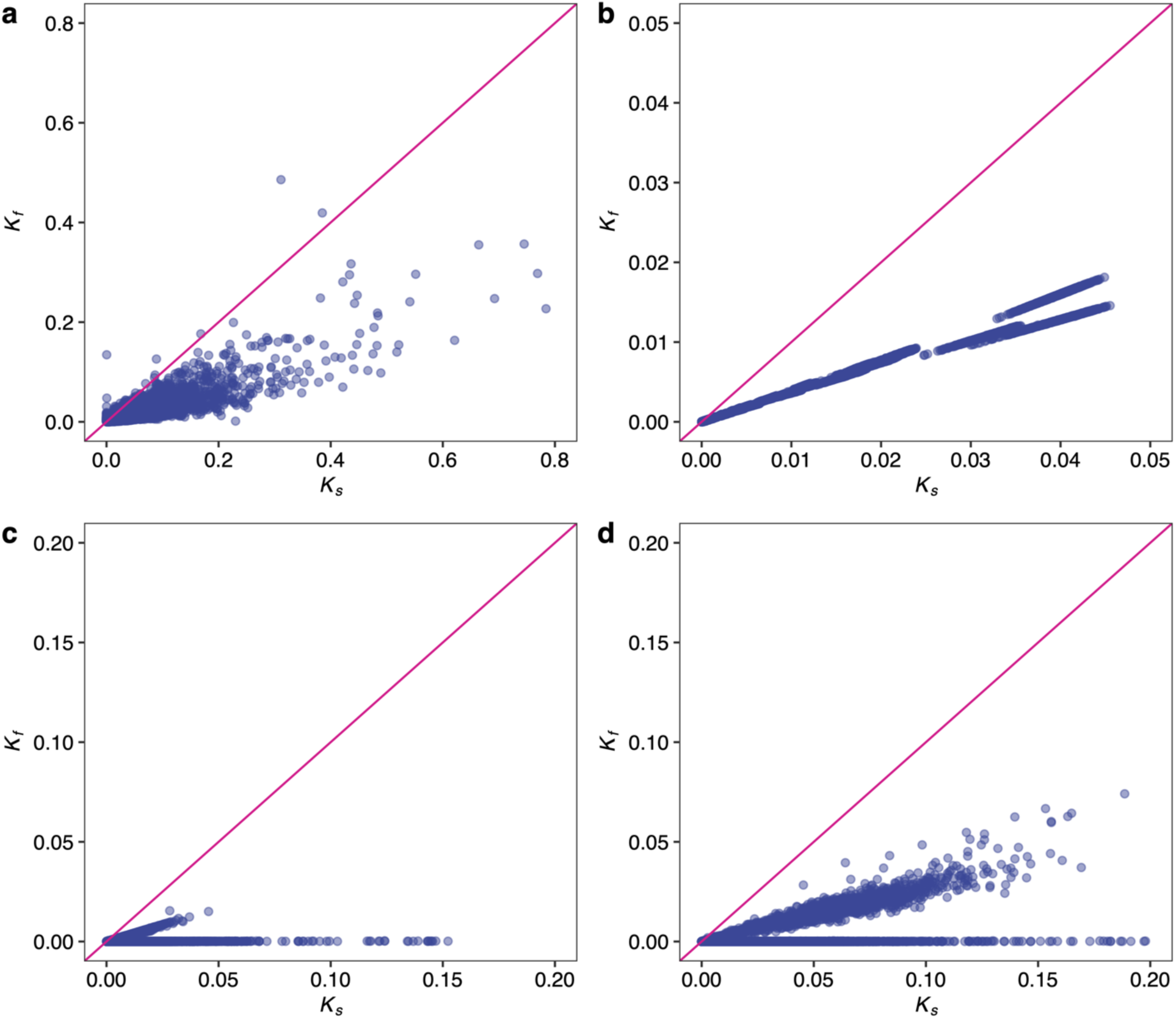
Relationship between structural keystones and functional keystoneness of each taxon over different samples. a, human gut microbiome. b, human oral microbiome. c, soil microbiome. d, coral microbiome.

